# The olfactory-based neurofeedback of the EEG alpha rhythm

**DOI:** 10.1101/2023.08.30.555545

**Authors:** Alexandra Medvedeva, Ivan Ninenko, Daria F Kleeva, Alexey Fedoseev, Artem Bazhenov, Miguel Altamirano Cabrera, Dzmitry Tsetserukou, Mikhail A Lebedev

## Abstract

Neurofeedback (NFB) is a form of biofeedback that enables subjects to monitor and control their own brain activity. To communicate with the subject, modern NFB training methodologies utilize various signaling mechanisms, typically visual and/or auditory stimuli. Olfaction has not been explored yet as a way to deliver NFB. Here we developed an olfactory-based NFB system based on electroencephalographic (EEG) recordings in human participants. The system incorporates an EEG recording apparatus, a custom olfactory display for the automated delivery of Sniffin’ Sticks, and a Python application for the conversion of EEG rhythms into NFB signals and controlling behavioral tasks. We tested occipital alpha rhythm as the source of NFB. Fifteen healthy participants were randomly assigned to three groups: olfactory neurofeedback, auditory NFB, and mock-olfactory NFB. NFB training resulted in an increase of alpha power in the true NFB groups, but not in the mock NFB group where the alpha power decreased, probably because of fatigue and drowsiness. Based on these results, we conclude that olfactory NFB is feasible, and lay out a framework for its future development.

## 1. Introduction

The neurofeedback (NFB) systems are a subtype of brain-computer interfaces (BCIs) where a user’s own neural activity is delivered back to the user in order to monitor and modify neural activity (Thibault, Lifshitz, & Raz, 2016). BCIs have proven their effectiveness as noninvasive tools for modulating neural activity, and have been integrated into numerous areas of human performance improvement and medical treatments of neurological conditions such as ADHD, epilepsy, depression and autism spectrum disorders (Marzbani, Marateb, & Mansourian, 2016, Thibault, Lifshitz, & Raz, 2016). NFB systems are closely related to brain-computer interfaces (BCIs), which use the same principles of operation but put a greater emphasis on the control of external devices with the user’s brain activity (Kawala-Sterniuk et al., 2021). The first NFB systems made use of the alpha rhythm, an EEG-derived signal occurring within the frequency range of 8-12 Hz. The alpha rhythm is one of the most robust brain EEG signals which is especially prominent in the occipital lobes. This rhythm is associated with the state of relaxed wakefulness, and it is particularly prominent over the occipital cortex when the eyes are closed. The characteristics of the alpha rhythm have been harnessed for therapeutic and cognitive enhancement purposes, with alpha training being one of the most well-studied and commonly used NFB protocols (Marzbani, Marateb, & Mansourian, 2016).

As an alternative to the predominant visual and auditory-based NFB, the present study probed an innovative approach that utilized olfactory NFB. The olfactory NFB was selected due to the unique sensory data it provides and the fact that it has been largely overlooked in previous NFB research. Recent studies on olfactory information processing demonstrate that the odor is encoded 100 ms after odor onset at the earliest, with its signal sources estimated in and around the olfactory areas. From the olfactory brain regions information spreads rapidly to the regions associated with emotional, semantic, and memory processing (Kato et al., 2022). It was demonstrated that pleasurable odors caused a testable increase in alpha rhythm power 8 s after odor delivery, but unpleasant sensations did not cause such an effect (Brauchli et al., 1995). Additionally, odor perception was shown to enhance EEG high-frequency components (11–25 Hz) with this effect being most pronounced in the occipital regions of the neocortex (Cherninskii et al., 2009). Interest in odor delivery techniques has been growing, and many types of olfactory display have emerged (Garcia-Ruiz, Kapralos, & Rebolledo-Mendez, 2021).

Here we integrated the historically effective alpha training NFB protocol with innovative olfactory feedback. Our aim was to explore new possibilities in NFB systems, potentially paving the way for more engaging, immersive, and effective NFB applications.

## 2. Materials and Equipment

### 2.1. EEG and respiration recordings

EEG data were collected with an NVX-52 amplifier (Medical Computer Systems, Russia) using three active electrodes placed over the parieto-occipital region: PO3, POz, PO4 (electrode names according to the International 10–20 system) and monopolar montage with a reference electrode at FCz. Data were recorded with a NeoRec computer program with a sampling rate of 500 Hz. A 0.5 Hz high pass filter, a 70 Hz low pass filter, and a 50 Hz notch filter were applied to the EEG signals.

### 2.2. Neurofeedback program

The neurofeedback program was written in the Python programming language using specialized software designed for EEG processing, MNE-Python (Gramfort et al., 2013). Frequency analysis was performed using Welch’s estimation of power spectra, where EEG signals were averaged within a 5-s time window in the frequency range of 1 - 30 Hz. Frequencies above this range were not considered in order to avoid unnecessary noise.

The threshold between successful and unsuccessful trials was adapted personally for each participant because the characteristics of the alpha rhythm are highly individual. The value of the threshold for training was calculated from the baseline recorded before the experiment as the ratio of the average power spectral density (PSD) in the frequency range of 8 - 12 Hz to the average PSD in the range of 1 - 30 Hz. While we used standard boundaries for the alpha-rhythm it is important to mention the concept of individual alpha frequency (IAF). IAF is a personalized alpha rhythm frequency that can slightly deviate from the standard alpha frequency band (8-12 Hz), due to factors such as age, mental state, arousal level, and even genetic characteristics (Klimesch, 1999). In this study we, however, chose to operate within the standard alpha frequency band of 8-12 Hz for the NFB protocols. This decision was made with the intention of ensuring comparable conditions across different participant groups and maintaining a uniform basis for comparison. Given that the alpha rhythm characteristics can differ considerably between individuals, utilizing a standardized frequency band helped to create a uniform approach for all participants, facilitating a more straightforward interpretation of the results.

The subjects received positive reinforcement if the value of this indicator exceeded the value of the threshold. The positive auditory neurofeedback consisted of a sequence of musical notes played through the headphones connected to the computer. The positive olfactory neurofeedback was provided with the computer-controlled olfactory display developed at the Skolkovo Institute of Science and Technology. The absence of a sound or odor was the negative feedback.

### 2.3. Group categorization algorithm

The participants whose data are reported here (N=15) were randomly assigned to three experimental groups: the olfactory, auditory, or control groups. The participants in the control group received a mock olfactory feedback. Each group consisted of five individuals. The categorization of participants was achieved via a computer program, ensuring randomness. Moreover, the study maintained a double-blind structure for the control and olfactory groups, facilitated by a software program used by the experimenter. This program determined the participant’s group assignment and initiated the relevant feedback protocol. It was only at the experiment’s completion that the experimenter was informed of the participant’s group affiliation.

### 2.4. Delivery of the olfactory stimuli

The olfactory stimuli were delivered by a computer-controlled olfactory display developed at the Skolkovo Institute of Science and Technology. This device is depicted in Figure 2. This custom-built device uses standard Sniffin’ Sticks, pens that are filled with fragrances instead of ink, which are widely used in olfactory research (Hummel et al., 1997). This robot for supplying odors consists of the following key electronic components: Arduino Mega as the main controller, a fan for direct airstream with the odor towards the subject, voltage regulators for power and logic power distribution, and main motors with precise positioning. This device carried out the supply of odor by raising from a sealed tube and holding an aromatic stick in front of a fan. Six sticks can be loaded into the robot and can be rotated to provide different odors. The last capacity has not been implemented in the current research as only the banana flavor stick has been presented during all trials. The smell of a banana was chosen as the most neutral in terms of severity, not as sharp as, for example, the smell of citrus, and according to subjective assessments of the participants, the smell of a banana did not cause sharp positive or negative sensations. The computer controlled the robot delivery and allowed the odor to be applied for a selected period, while the rest of the time the odor was completely absent. The current prototype required 1 second to open the tube and raise the stick to the point where the participant would be able to perceive the odor.

**Figure 2.**
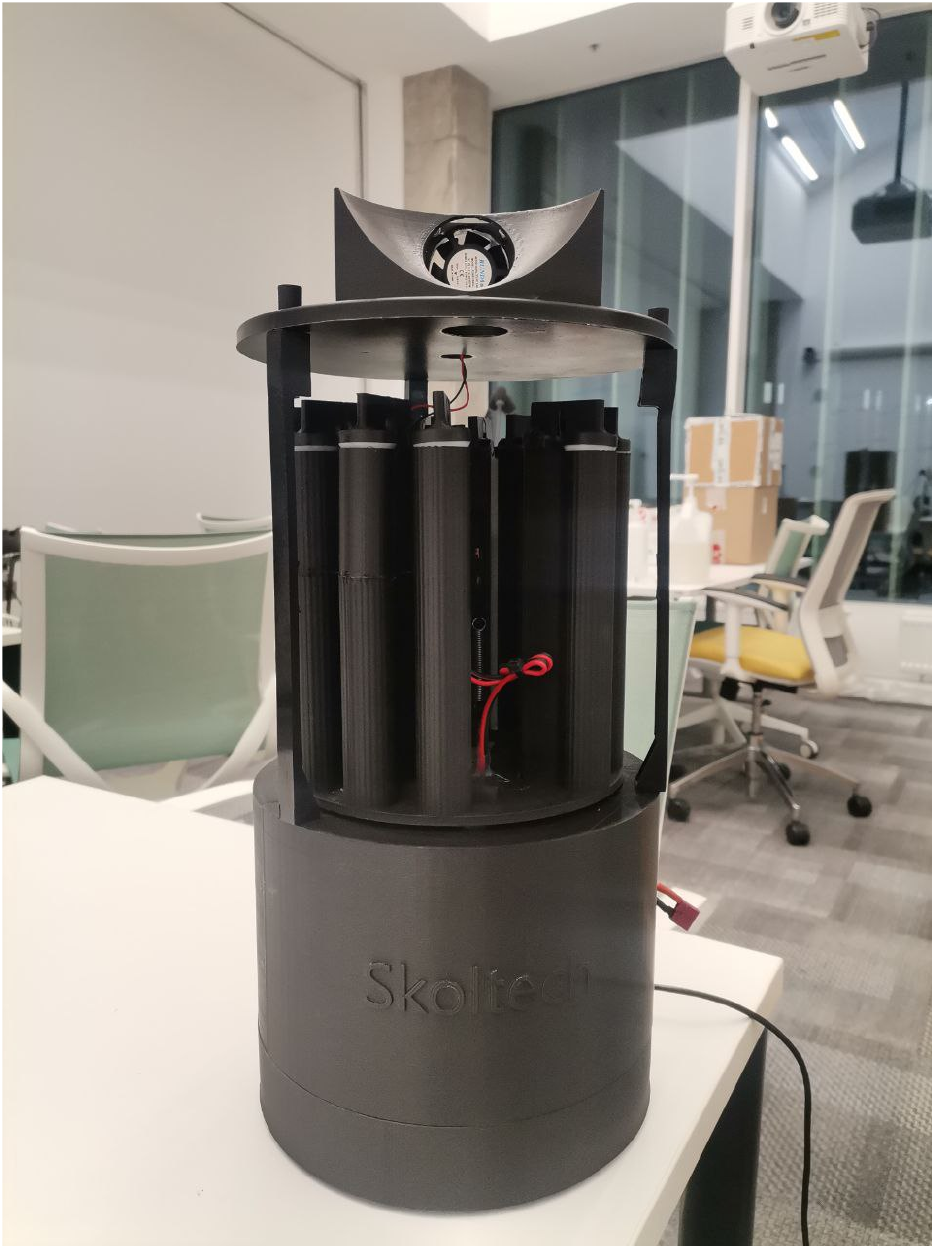
A photograph of the olfactory display used in the study.

## 3. Methods

### 3.1. Subjects

The experiment was carried out at the Vladimir Zelmen Skoltech Center for Neurobiology and Brain Rehabilitation. Healthy volunteers aged 18 to 35 were invited to participate in the experiment through an open call. The Ethics Research Committee of the Skolkovo Institute of Science and Technology (Skoltech) approved the experimental protocol of this study. All participants provided written informed consent prior to the experiments.

Participants did not receive any financial reward for this experiment. All of them were familiarized with the experimental procedure and asked to fill a form after the experiment. The form included general sociodemographic information and some specific questions regarding olfaction. This form was adapted from the previous research on olfactory stimuli (Ninenko et al., 2023).

A total of 17 participants (10 males and 7 females, one left-handed, all others right-handed, aged 24.29 ± 2.4 years, mean ± SD) took part in the experiment. Due to problems with the signal quality, two of them were removed from the final sample. We report the results from 15 participants (9 males and 8 females, one is left-handed, all others are right-handed, aged 24.00 ± 2.2 years, mean ± SD). Participants were randomly divided into three experimental groups: olfactory or auditory, and the control group which received mock olfactory feedback. The distribution of participants into groups was randomized using a computer program and the study was double-blind, as the experimenter ran a program that determined the subject’s belonging to one of the groups and launched the necessary feedback protocol.

### 3.2. Study design

The experimental design was crafted with the aim of examining both olfactory and auditory feedback in conditions that were as identical as feasible. This required overcoming a notable constraint tied to olfaction - the fact that humans detect odors during inhalation, but not during exhalation. (This can be likened to the process of blinking, which could potentially delay the reception of visual stimuli.) In the context of olfaction, the time spent exhaling equals or even exceeds the time taken to inhale. This particularity restricts the ability to generate real-time feedback in the same manner as it would be done for visual or auditory feedback.

It would indeed be possible to construct an experiment that factors in the inhalation cycle when delivering feedback, but such an arrangement would not be readily adaptable to other sensory inputs. To circumvent these constraints, a delayed-feedback design was selected. While this could have marginally compromised the efficiency of the alpha training, this design facilitated a test of olfaction and auditory feedback under conditions that were closely matched. Auditory feedback was preferred over visual feedback, given its ability to be perceived even with the eyes closed. This refined experimental design thereby ensured that the only variable element was the sensory channel delivering the feedback, with all other factors kept constant.

### 3.3. Behavioral task

Prior to initiating the experiment, participants were thoroughly briefed regarding the structure of the experiment and were encouraged to aim for maximum positive reinforcement. They were informed about their group assignment, whether it was auditory or olfactory feedback. No guidance was provided in terms of employing a specific cognitive strategy. While participants were made aware of the possibility of receiving sham feedback, neither the participant nor the experimenter were aware if the participants were allocated to the control group.

The experiment began by recording a 40-s baseline with eyes closed. The eyes remained closed throughout the subsequent session. Based on the baseline, the threshold value was determined. The training period consisted of a sequence of 60 sessions, each consisting of a 10-s trial and a 10-s feedback session, being 20 minutes in total. NFB training took 2 days. Thus, each subject took part in two similar pieces of training on two consecutive days, around the same time of day.

During the experimental session, EEG data were continuously sampled and the alpha rhythm was compared to the baseline. After each 10-s trial, the participant received a sound signal in the headphones (a tune), announcing the start of the feedback. This was followed by a 10-s NFB session. The end of the NFB session was also announced by a tune. Those tunes remained the same for the duration of the experiment. In the auditory group, participants were instructed to pay attention to the presence of tunes in between the starting and the ending tunes. In the case of positive NFB, a series of tunes was played. In the olfactory group, participants were instructed to pay attention to the presence of odor during the NFB session. In the case of positive NFB, an odor was delivered by the olfactory display. The scheme of the experiment is shown in Figure 3.

**Figure 3.**
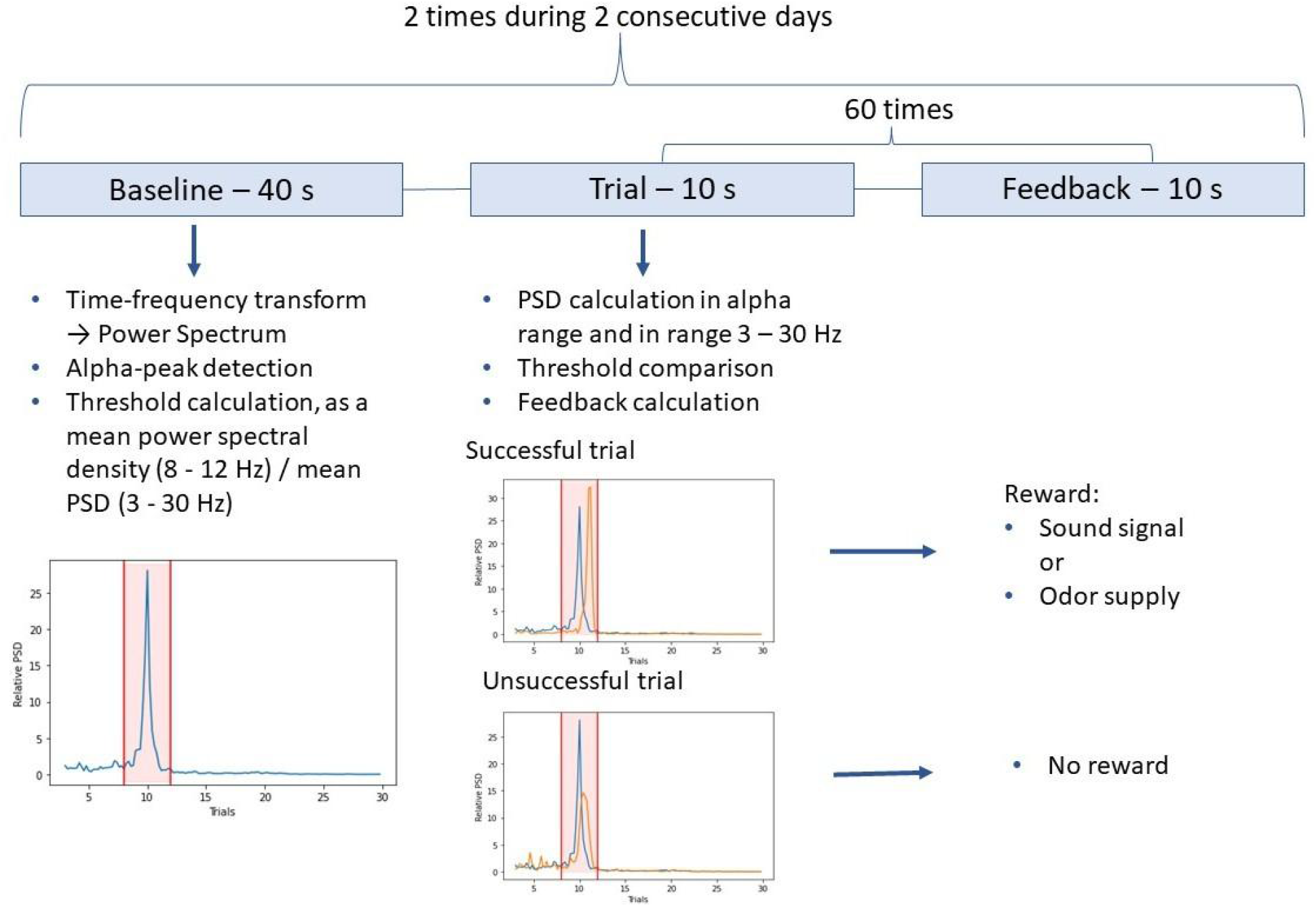
A schematic representation of the experimental sequence.

The experimental procedure was executed to ensure consistent conditions across all participant groups, ruling out the influence of any extraneous variables. Throughout the entirety of the experiment, a steady air stream was maintained toward the participant’s face, delivered via an odor display fan. This setup ensured no perceptible variation in the experimental conditions, effectively eliminating any potential confounding effects arising from external sound, visual cues, or tactile stimuli produced by the odor display fan.

In order to further control for external sensory stimuli, the experiments were conducted in a dimly lit room. This subdued lighting, coupled with the participants’ eyes being closed throughout the experiment, served to heighten the focus on the feedback mechanisms being studied. To cater to the different feedback modalities used in the experiment, participants were fitted with headphones. For the control and olfactory groups, these headphones provided the necessary signals indicating the commencement and conclusion of the feedback sessions. For the auditory group, the headphones were crucial for the delivery of the feedback itself. Further, to minimize any potential noise from the odor display or any other extraneous sources, soundproof headphones were used. This helped in ensuring that the auditory environment was strictly controlled, providing a stable sensory backdrop against which the effects of the different feedback mechanisms could be accurately assessed.

### 3.4. Data processing

The data were processed using the Python programming language and specialized software designed for EEG processing, called MNE-Python (Gramfort et al., 2013).

When calculating the power spectral density, the same parameters were used as for calculating the threshold during the experiment. Frequency analysis was performed using Welch’s estimation of power spectra, followed by averaging across the electrodes with a time window of 5 s in the frequency range of 1 - 30 Hz. To average the data of the participants, a z-score was calculated, which improved the analysis of the individual differences in the alpha PSD of the participants and subsequently compared the average indicators across the groups. To highlight the main trends in alpha PSD over the sessions, a rolling average was calculated with a window covering 20 trials.

For an additional estimate of the alpha PSD, the signal envelope was calculated. The envelope spectrum estimated the periodicity of all band-limited activity. To calculate the envelope, the signal was filtered in the frequency band 8 - 12 Hz, and then the temporal envelope was extracted using the Hilbert transform (Emmert et al., 2017).

## 4. Results

### 4.1. Individual results from the training sessions

The percentage of successful attempts in the sound-feedback group was 3-48% (20%± 19.5%, mean ± SD) on the first day and 3-50% (24% ± 18.2%, mean ± SD) on the second day. In the olfactory feedback group, the percentage of successful attempts was 21-90% (47%± 25.9%, mean ± SD) on the first day and 10-88% (43% ± 31.7%, mean ± SD) on the second day. These differences were associated with the individual characteristics of the subjects.

Figure 4 shows the change in the relative PSD of the alpha rhythm for one of the participants from the olfactory group. Both experimental days are shown. On the first day, there was a decrease in the alpha-range PSD. On the second day, the alpha-range PSD was near-constant level, and the number of successful attempts increased as compared to the first day of research.

**Figure 4.**
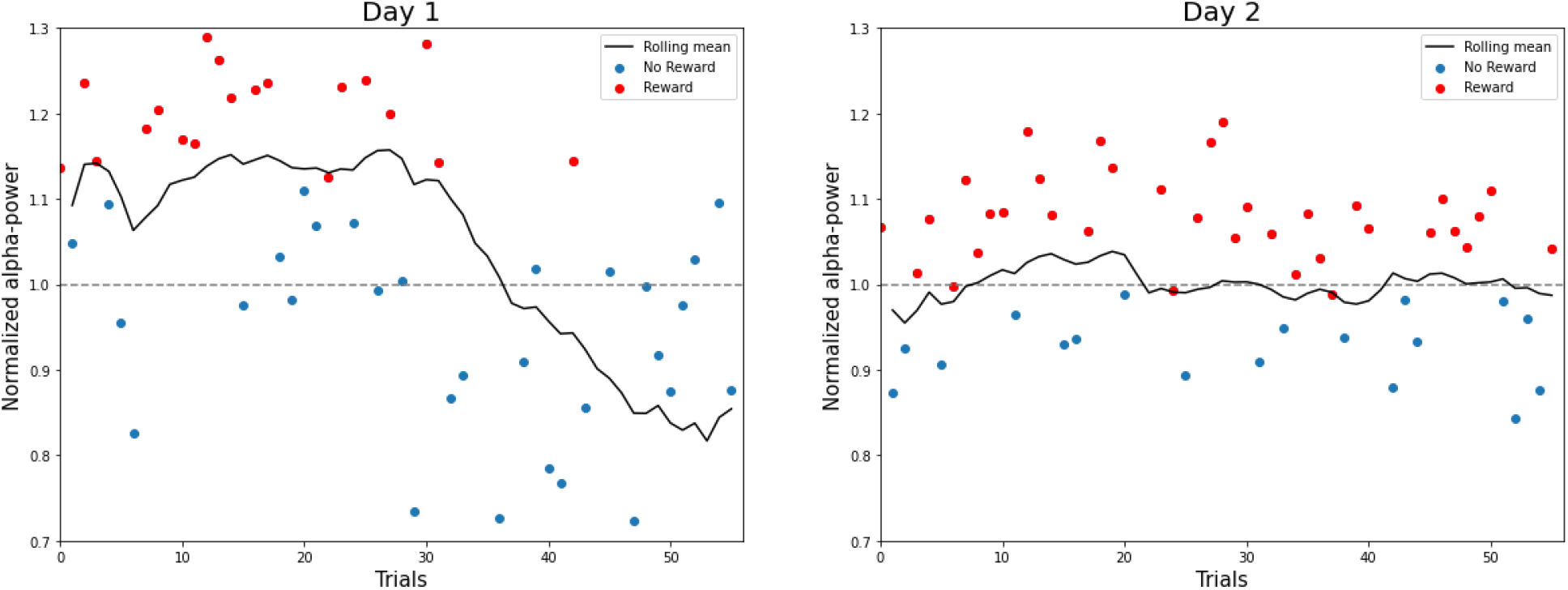
The change in the relative PSD of the alpha range for one of the subjects from the olfactory group during the training sessions. Data from both training days are shown. The dots indicate PSD for each trial normalized for the average value of the PSD per session, the colors correspond to the attempts that were followed or not followed by a reward. The black curve is the rolling mean (with a 20-point window).

Figure 5 shows the EEG power spectra of one of the participants in the olfactory group on the first day of the experiment. An alpha peak stands out in the spectrum. When calculating the PSD of the alpha rhythm, a standard frequency range of 8-12 Hz was utilized (highlighted in green in Fig. 5). However, it was observed that the alpha peak did not always align perfectly within this range in all people. Therefore, for future studies, we would recommend that an individualized alpha range is employed and calculated specifically for each participant.

**Figure 5.**
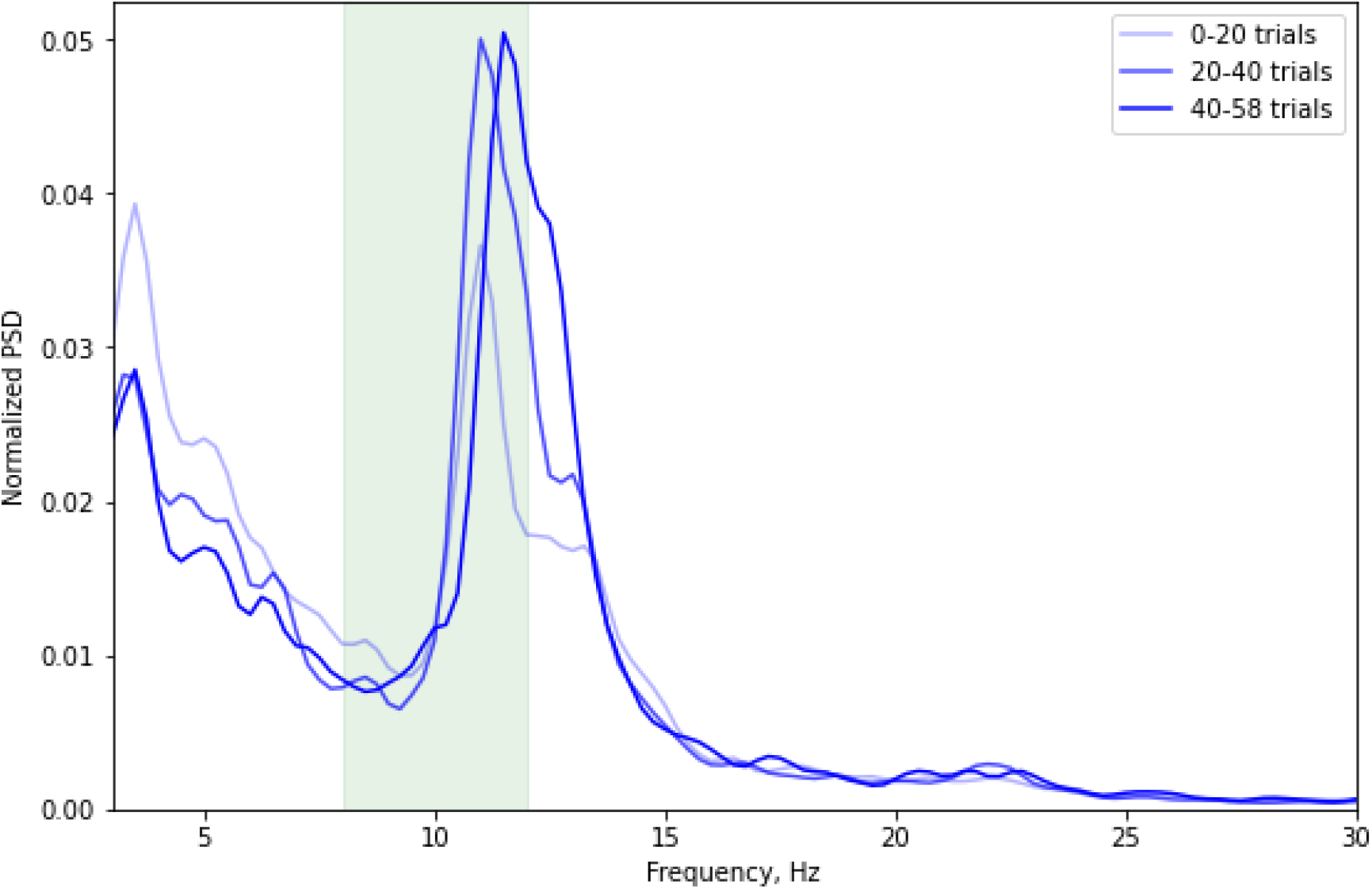
EEG power spectra of a subject from the olfactory group. The spectra for the first, second, and third thirds of the experimental session are shown with different line color. The green rectangle highlights the alpha range (8 – 12 Hz).

Changes in the alpha power are visible at different time intervals of the session. There is an increase in the alpha peak from the first to the last third of the experiment, as well as a decrease in power in the low-frequency range.

### 4.3. Alpha-range power changes

Figure 6 shows the change in the moving average (with a 20-point window) of the PSD of the alpha range during the experimental sessions, normalized to the power of the first 10 attempts. In the control group, on both days, a gradual decrease in PSD was observed. On the second day the decrease occurred almost immediately, and continued until the end of the experiment. In the experimental groups, there was a slight increase in PSD followed by a decrease almost to the initial level. Except for the olfactory group on the first day, in the experimental groups a gradual decrease in power occurred around the 20th attempt of the session, which corresponded to approximately minute 7 of the experiment. Probably, by this time, many subjects experienced fatigue associated with excessive concentration on the task and/or drowsiness associated with a long stay in a static position with their eyes closed. On the first day of the experiment, the decrease in the PSD remained constant until the end of the experiment. On the second day, the decrease was followed by an increase.

**Figure 6.**
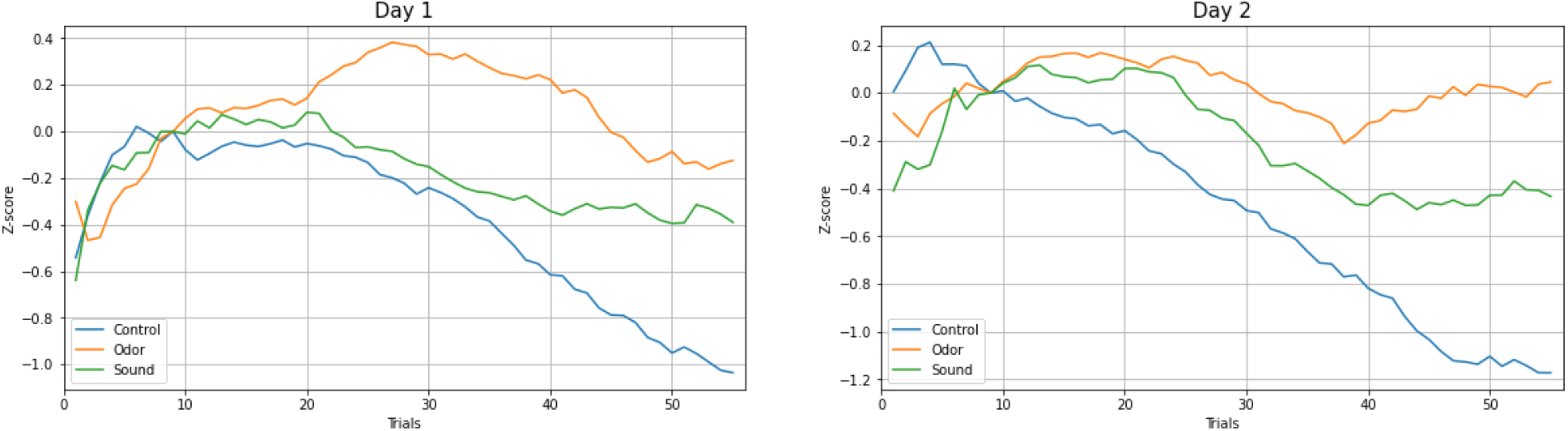
The change in the rolling mean from the z-score of alpha-range PSD averaged over the group of subjects, during the experimental sessions of two days, normalized to the first 10 trials for more visual clarity. Experimental groups are shown in different colors.

Similar results were obtained when calculating alpha PSD using signal envelope (Fig. 7). In this case, there was also a decline in PSD for the control group. At the same time, there was no pronounced decrease in power in the auditory and olfactory groups.

**Figure 7.**
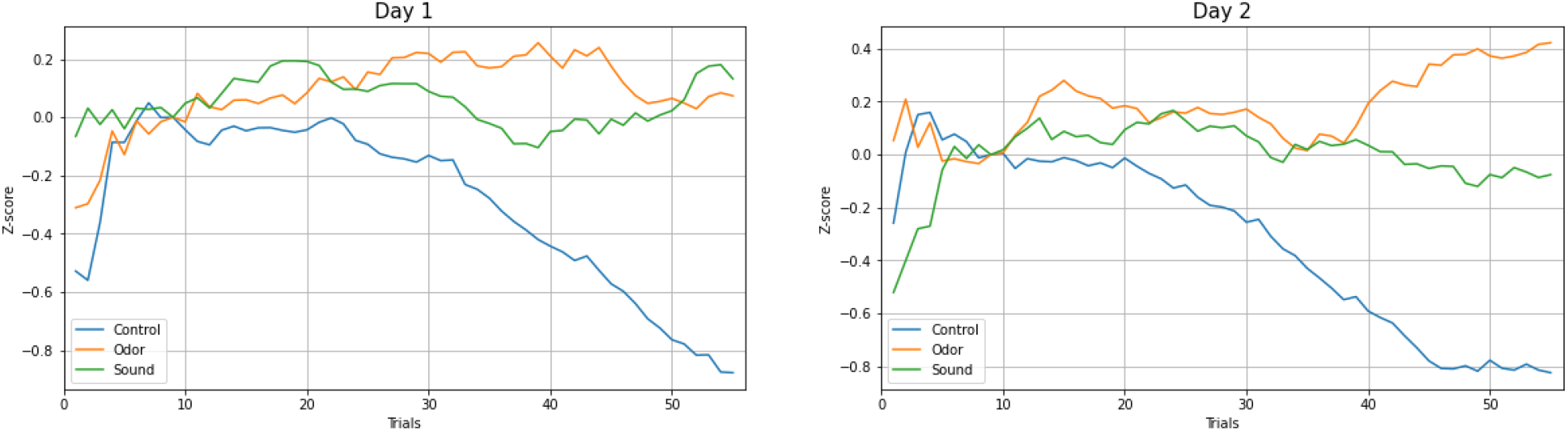
The change in the rolling mean from the z-score of alpha-range envelope PSD averaged over the group of subjects, during two daily experimental sessions, normalized to the first 10 trials. The experimental groups are indicated by color.

Thus, in this experiment, NFB contributed to the maintenance of the average level of PSD in the alpha range in the experimental groups, which was evident from the comparison to the control group.

The literature reports poor results with neurofeedback having a long time delay. Thus, continuous feedback led to a faster control compared to intermittent feedback in the studies with visual NFB derived from fMRI (Oblak, Lewis-Peacock, & Sulzer, 2017, Schütze & Junghanns, 2015). Additionally, sleep was reported to easily occur during alpha/theta NFB training (Le Van Quyen et al., 2001). Many of our subjects reported high workload and sleepiness, which have also been shown to reduce the power of the alpha rhythm [16]. The fact that the subjects had to keep their eyes closed throughout the experiment also could have had an effect, which could have contributed to increased drowsiness. In the future, it would be worth reducing session time to 10 min and increasing the number of training sessions.

## 5. Discussion

In this study, we probed the feasibility of an olfactory NFB. The experiments were designed to allow comparison between the olfactory and auditory feedback. An important factor related to olfaction, where odors are perceived during inhalation but not exhalation, was incorporated in the experimental paradigm. The experiment employed a delayed-feedback which maintained comparable conditions across the sensory modalities.

We observed an increase in the relative PSD in the alpha-range during the first 20 trials (7 minutes) of the experimental session for the participants receiving NFB. On the first day of training, a decrease in alpha power occurred next, and the power remained at a near-constant level until the end of the session. On the second day, the sequence of alpha changes was similar to the first day, with a difference that the power slightly increased towards the end of the session. Similar patterns of EEG changes were observed for the olfactory and auditory feedback. By contrast, for the control group, a gradual decrease in the alpha PSD was observed during both experimental days. Thus, for the NFB settings employed, the olfactory feedback was as effective as the auditory feedback. The absence of any dramatic increase in the alpha power for both the olfactory and auditory NFB could be related to the fact that thefeedback arrived late and infrequently.

The future advancement of olfactory NFB should be based on the improved NFB settings. These settings should be optimized for olfaction rather than copying the designs for the other sensory modalities. Central to this approach is the integration of the breath cycle into the NFB design. Incorporating the measurements of respiration acknowledges the inherent physiological process of odor perception occurring predominantly during inhalation. The proper NFB design should assist the user in achieving a comfortable and appropriate breathing rhythm. Such breath cycles would be associated with the brain states that need to be modified by NFB. Here we showed it is feasible to modulate the alpha rhythm with olfactory NFB. The capacity of such a NFB to modulate the other EEG rhythms should be addressed by future research. Further research and optimization in this realm will pave way for novel, efficient, and effective therapeutic and training solutions.

